# Prophylaxis and Treatment of SARS-CoV-2 infection by an ACE2 Receptor Decoy

**DOI:** 10.1101/2022.12.31.522401

**Authors:** Takuya Tada, Belinda M. Dcosta, Hao Zhou, Nathaniel R. Landau

## Abstract

The emergence of SARS-CoV-2 variants with highly mutated spike proteins has presented an obstacle to the use of monoclonal antibodies for the prevention and treatment of SARS-CoV-2 infection. We show that a high affinity receptor decoy protein in which a modified ACE2 ectodomain is fused to a single domain of an immunoglobulin heavy chain Fc region dramatically suppressed virus loads in mice upon challenge with a high dose of parental SARS-CoV-2 or Omicron variants. The decoy also potently suppressed virus replication when administered shortly post-infection. The decoy approach offers protection against the current viral variants and, potentially, against SARS-CoV-2 variants that may emerge with the continued evolution of the spike protein or novel viruses that use ACE2 for virus entry.

## Introduction

Since the initial zoonosis of SARS-CoV-2 into humans, the virus has undergone rapid evolutionary adaptation to the new host with the appearance of variants with divergent spike proteins. The appearance of the earlier Variants of Concern (Alpha, Beta, Gamma and Delta), each with a few mutations in the receptor binding domain, were replaced by the sudden emergence of the highly divergent Omicron variant first reported in Botswana and South Africa (CDC, 2022). As the virus has continued to adapt to its human host, it has become increasingly transmissible as a result of mutations that increase the affinity of the spike protein for the ACE2 receptor. Omicron BA.1 rapidly became prevalent world-wide and then gave rise to the BA.2 subvariant (CDC, 2022) and more recently to the highly transmissible BA.2.12.1, BA.4, BA.5, BA.2.75 and XBB subvariants, some of which have increased affinity for ACE2 (Cao et al., 2022b; Yue et al., 2023).

The divergence of the spike protein has presented an obstacle both to the effectiveness of vaccines and to monoclonal antibody therapy. The Regeneron REGN-COV2 monoclonal antibody cocktail and Lilly monoclonal antibodies that had potent neutralizing activity against earlier Variants of Concern spikes (Baum et al., 2020; Chen et al., 2021a; Chen et al., 2021b; Hansel et al., 2010; Planas et al., 2021b; Tada et al., 2021a; Tada et al., 2022b; Tada et al., 2021b; Tada et al., 2022c; Wang et al., 2021; Weinreich et al., 2021; Weisblum et al., 2020) and had been highly effective at preventing hospitalization and morbidity of patients infected by the earlier Variants of Concern, were rendered ineffective by the heavily mutated spikes of the Omicron variants which escape neutralization (Cameroni et al., 2021; Cao et al., 2021; Hoffmann et al., 2021; Iketani et al., 2022; Liu et al., 2022; Planas et al., 2021a; Tada et al., 2022a; VanBlargan et al., 2022; Zhou et al., 2022). The Omicron variants pose an additional obstacle to the prophylactic use of monoclonal antibodies (Cameroni et al., 2021; Cao et al., 2021; Hoffmann et al., 2021; Iketani et al., 2022; Liu et al., 2022; Planas et al., 2021a; Tada et al., 2022a; VanBlargan et al., 2022; Zhou et al., 2022). The AstraZeneca dual monoclonal antibody cocktail Evusheld is used primarily for prophylaxis in immunocompromised individuals for whom vaccination may be ineffective (ClinicalTrials.gov,); however, both Evusheld antibodies have significantly decreased neutralizing titers against the BA.1 and BA.2 variants which could affect the long-term effectiveness of the therapy (Cao et al., 2021; Iketani et al., 2022; Liu et al., 2022; Planas et al., 2021a; Tada et al., 2022a; Zhou et al., 2022). The Sotrovimab monoclonal antibody Vir-7831 retains activity against the earlier Variants of Concern but has substantially decreased neutralizing titer against the BA.1 and BA.2 variants. Until recently, the only therapeutic monoclonal antibody that neutralized the Omicron variants was LY-CoV1404 (Westendorf et al., 2022); however, recent viral subvariants escape neutralization by the monoclonal antibody. In light of the continued evolution of viral variants with mutated spike proteins, there is a need for improved treatment and therapies that are less affected by spike protein variability.

The concept of decoy receptors for the treatment of virus infection was initially tried as a therapy for HIV infection. Decoys are predicted to be less susceptible to escape by mutagenesis of the viral spike protein and less like to induce an antibody response as they are derived from self-protein sequences to which the immune system is tolerant. A soluble CD4 “immunoadhesin”, in which the ectodomain of CD4 was fused to an immunoglobulin domain was previously reported (Traunecker et al., 1989). Although the protein neutralized the virus by binding the gp120 subunit of the viral envelope glycoprotein *in vitro* (Daar et al., 1990; Haim et al., 2009; Orloff et al., 1993; Schenten et al., 1999; Sullivan et al., 1998), it failed to decrease virus loads when used to treat patients. More recently the concept was revived using an adeno-associated virus vector expressing an enhanced soluble eCD4-Ig decoy (Gardner et al., 2015; Spitsin et al., 2020). Rhesus macaques treated with the vector were highly protected against a challenge with SHIV-AD8 and SIVmac239 (Gardner et al., 2015; Spitsin et al., 2020).

We previously reported on a receptor decoy protein for SARS-CoV-2 termed a “microbody” in which the ACE2 ectodomain is fused to the CH3 domain of an immunoglobulin IgG1 heavy chain (Tada et al., 2020). Truncation of the Fc domain served to decrease the mass of the protein as well as to prevent binding of the protein to cell surface Fcγ receptors (Maute et al., 2015). The ACE2 ectodomain contained a point mutation (H345A) that inactivates the carboxypeptidase activity of ACE2 (Guy et al., 2005) to prevent possible effects of the protein on blood pressure. The protein potently neutralized the parental D614G virus and viruses with the variant of concern spike proteins by binding to the viral spike protein, preventing association of the virus with cell surface ACE2 (Tada et al., 2020). Because ACE2 binding is a conserved feature of all SARS-CoV-2 spike proteins, the decoy is predicted to maintain its neutralizing activity against current, as well as future virus variants, without being affected by mutations in the variant spike proteins.

In this report, we tested a decoy protein mutated to increase its affinity for the spike protein containing a truncated Fc region for its ability to prevent and treat SARS-CoV-2 infection in mouse models. We found that the recombinant protein was a potent prophylactic against both parental SARS-CoV-2 and the Omicron variants and was an effective therapeutic that rapidly decreased virus loads when administered post-infection. The findings confirm and extend findings from other groups using decoy.Fc fusion proteins (Higuchi et al., 2021; Ikemura et al., 2022; Zhang et al., 2022).

## Results

### High affinity decoy inhibits SARS-CoV-2 infection and replication *in vitro*

We previously reported the construction of plasmid vectors expressing an ACE2 microbody (pcACE2.mb) in which the ectodomain of human ACE2 is fused to the IgG1 CH3 domain and soluble ACE2 (pcsACE2) that encodes the unfused ectodomain (Tada et al., 2020). To further increase the potency of the microbody decoy protein, we induced the mutations T27Y/L79T/N330Y reported by Chan *et al*. (Chan et al., 2020) that increase ACE2 affinity for the SARS-CoV-2 spike protein (pcACE2.1mb) (**Figure 1A**). In addition, the proteins are mutated in the catalytic active site at position 345 (H345A) to inactivate phosphohydrolase activity (Guy et al., 2005), preventing possible effects on blood pressure, and contain a carboxy-terminal His-Tag (**Figure 1A and 1B**). The decoy proteins were produced by transfection of ExpiCHO cells with the expression vectors and then purified by affinity chromatography and size exclusion chromatography (**Figure 1C**). The antiviral activity of the decoy proteins was tested in the pseudotyped lentivirus neutralization assay. The results show that the ACE2.mb decoy neutralized virus with the D614 spike protein with a potency 10-fold increase compared to sACE2 while the high affinity ACE2.1mb decoy increased neutralizing activity another 5-fold (**Figure 1D**). The decoy neutralized virus with the D614G spike with a similar potency. The ACE2.1mb decoy was also active against virus with the Alpha, Beta, Gamma and Delta spike proteins. Analysis of decoy antiviral activity against live virus showed that the decoy suppressed the replication of USA-WA1/2020 and the Omicron variants. The ACE2.1mb decoy was 35-fold and 4-fold more potent than sACE2 and the ACE2.mb, respectively (**Figure 1E**). The decoys were active against BA.1 and BA.2 although the variants showed a degree of resistance with a 6.2-fold and 16-fold decreased titer, respectively.

**Figure 1.**
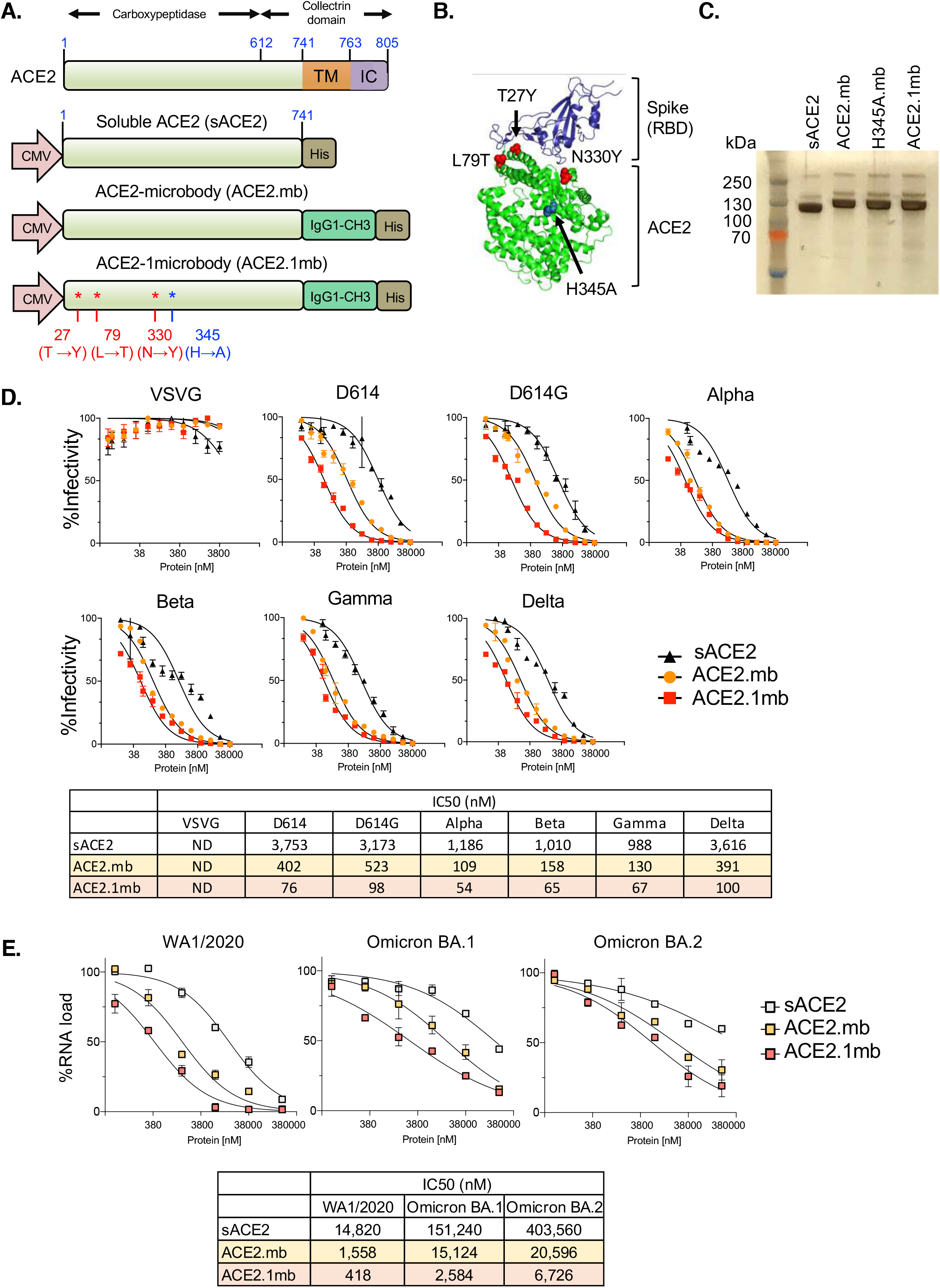
High affinity ACE2 decoy potently neutralizes SARS-CoV-2 variants. (A) The domain structure of soluble ACE2, ACE2mb and high affinity ACE2.1mb is shown. The ectodomain is shaded green; transmembrane domain (TM), intracellular domain (IC), the IgG1 CH3 domain is indicated and carboxy-terminal 6XHis-tag are shown. The high affinity ACE2.1mb decoy has mutations to increase spike protein affinity (T27Y, L79T, N330Y) and activate site point mutation H345A to inactivate carboxypeptidase activity. The genes are cloned into expression vector pcDNA3.1 in which transcription is driven by the cytomegalovirus promoter. (B) The 3D structure of the ACE2 (green) and spike protein (purple) complex was generated using PyMoL. The position of the mutations for improved decoy in the ACE2 carboxypeptidase domain are shown. (C) Recombinant decoy proteins produced in transfected ExpiCHO cells were purified by Ni-NTA affinity chromatography followed by size-exclusion chromatography. Purity of the proteins (25μg) were analyzed on a silver staining. (D) Neutralizing activity of the decoy proteins was measured using the variant spike protein-pseudotyped lentiviruses assay with a packaged luciferase expressing lentiviral genome. Lentiviruses were pseudotyped by the ancestral spike, parental spike or Alpha, Beta, Gamma and Delta spike protein. Vesicular stomatitis virus G protein (VSV-G) pseudotype served as a specificity control. Variable amounts of decoy (indicated on the X-axis) were incubated with a fixed amount of pseudotyped lentivirus (MOI=0.2). Infectivity (indicated on the Y-axis) is displayed as the percent infection normalized to control untreated virus as determined by luciferase assay of the infected cultures. The IC50s (nM) of decoy neutralization on each variant spike protein pseudotyped virus is calculated using the neutralization curves is shown in the table below. (E) 2-fold serial dilutions of decoy proteins were incubated with USA-WA1/2020, Omicron BA.1 or BA.2 virus (MOI=0.01) and added to Vero E6. After 2 days of infection, cells were harvested and subgenomic E gene was quantified by RT-PCR. IC50s (nM) were calculated from curves using Prism GraphPad 8 software and shown in the table. The experiment was done three times with similar results.

Avidity of the decoys for the spike protein was analyzed in two assays (Tada et al., 2020; Tada et al., 2022b). The first was a virion binding assay that measured the binding of spike protein-pseudotyped virions to bead-bound decoy protein. The results showed that the ACE2.1mb decoy bound more avidly to virions than sACE2 or the ACE2.mb (**Supplementary Figure 1A**). The second assay measured the binding of sACE2-nLuc, ACE2.mb-nLuc, ACE2.1mb-nLuc decoy:nanoluciferase fusion proteins to bind to spike protein-expressing 293T cells (**Supplementary Figure 1B and C**). The results confirmed the increased avidity of ACE2.1mb as compared to sACE2 and ACE2.mb for spike protein binding and showed the increased potency of the decoy for the Alpha, Beta, Gamma and Delta spikes and a 4- and 5.9-fold decrease in binding to BA.1 and BA.2 spike proteins (**Supplementary Figure 1D**). Taken together, the results showed that the ACE2.1mb decoy protein was a potent inhibitor of parental and variant SARS-CoV-2.

### Increased half-life of microbody decoy *in vivo*

Fusion of Fc domains onto proteins has been used to increase their half-life *in vivo* (Czajkowsky et al., 2012; Roopenian and Akilesh, 2007). To determine whether the truncated single IgG1 CH3 domain of the microbody decoy would extend its half-life and to determine the tissue localization of the decoy, recombinant decoy proteins sACE2-nLuc and ACE2.1mb-nLuc were produced. The fusion proteins were injected intraperitoneally (i.p.) and the mice were live-imaged over 3 days. The results showed that the proteins localized mainly to the spleen and that the sACE2 decoy was nearly undetectable after one day, the ACE2.1mb protein was detectable after 3 days (**Figure 2A**). To further analyze the tissue distribution of the ACE2.1mb-nLuc decoy and to understand how the route of administration affects its distribution, The ACE2.1mb decoy was injected i.p. or intravenously (i.v.) or instilled intranasally (i.n.) and 72 hrs post-administration, the amount of decoy protein in different tissues was determined by measuring the luciferase activity in lysates of individual organs. In mice injected i.p, the decoy localized mainly to the spleen, liver and serum with a minor fraction in the lung. I.v. injection similarly localized the decoy to the spleen, liver and serum but resulted in a 100-fold increase in localization to the lung. I.n. instillation resulted in localization of the decoy to the lung and trachea, as might be expected, with a small amount of the protein in the nasal tissues and serum (**Figure 2B**). To determine the half-life of the decoy *in vivo*, the ACE2.1mb decoy was administered i.v. or i.n. and luciferase activity in the serum and lung was measured over 20 days. The half-life of the i.v. injected or i.n. instilled decoy in the serum was 5.2 and 5.1 days, respectively. In the lung, the half-life of the i.v. injected protein was 4.0 days and i.n. instilled protein was 7.6 days (**Figure 2C**). The increased half-life and localization to the lung by the i.n. instilled protein suggests that this route of administration would be most effective therapeutically while i.v. injection would also be effective.

**Figure 2.**
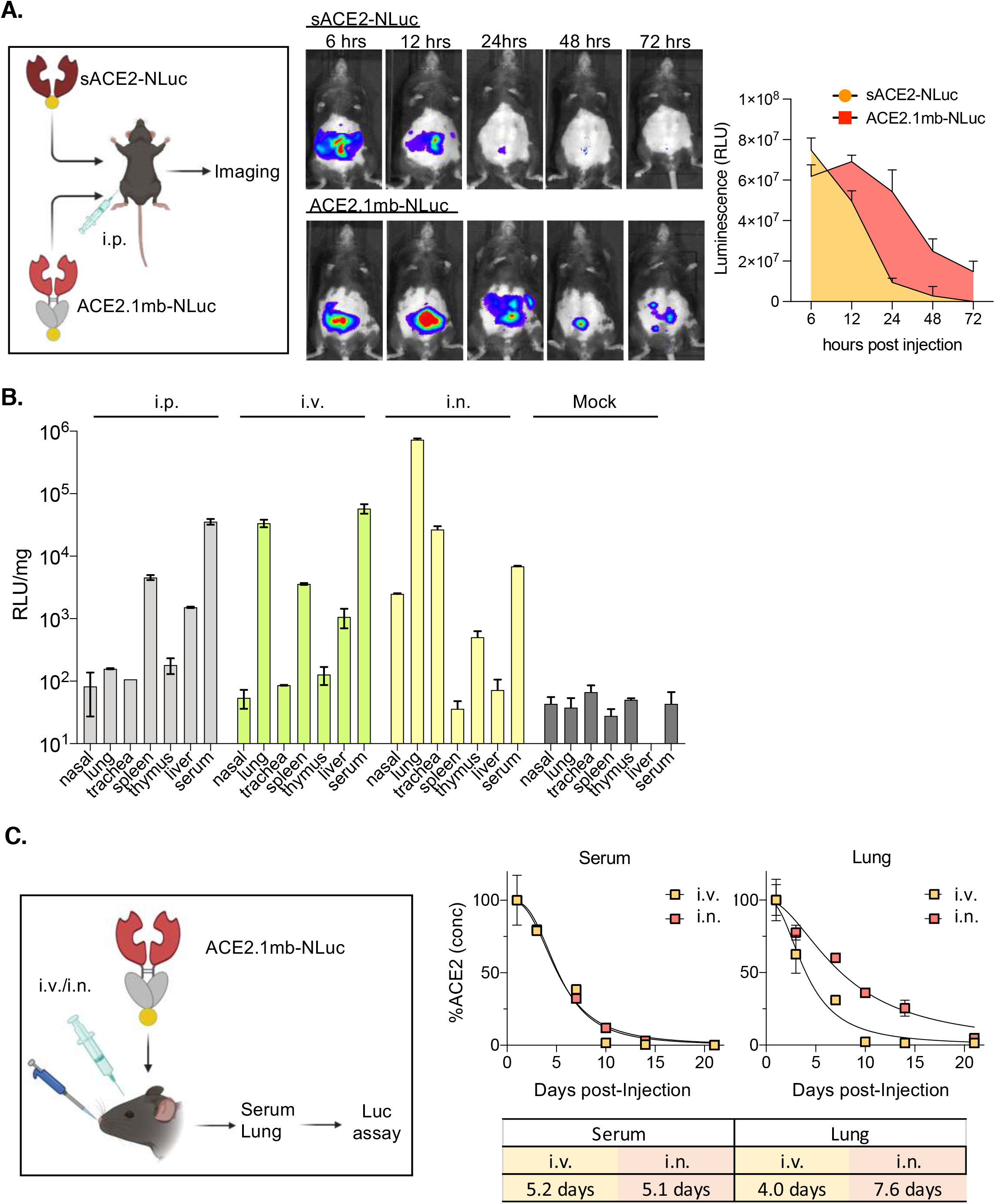
ACE2.1mb is stable *in vivo*. C57BL/6 mice were injected i.p. with 100 μg of sACE2.nLuc or ACE2.1mb.nLuc nanoluciferase fusion proteins. After 6, 12, 24, 48 and 72 hours of injection, the mice were imaged with Nano-Glo substrate. The panel on the right shows a graph of the fluorescence in relative light units (RLU). (B) ACE2.1mb-nLuc (100 μg) was administered i.p., i.v. or i.n. to C57/BL6 mice (n=3). 3-dpi, nasal epithelium, lung, trachea, spleen, thymus, liver and serum were harvested and luciferase activity was quantified. (C) ACE2.1mb-nLuc protein (100 μg) was administered by i.v. injection or i.n. installation. After 0.25, 1, 7, 10, 21 days, lungs and serum were harvested and luciferase activity was quantified. The concentration and half-life of decoy was determined based on standard curve obtained from the mixture of serial diluted ACE2.1mb-nLuc proteins and tissue lysate from wild-type (n=3). The experiment was done twice with similar results.

### Decoy protects against SARS-CoV-2 infection

To test the ability of the high affinity decoy to prevent SARS-CoV-2 infection *in vivo*, hACE2KI (Knockin) mice that have a knock-in of human ACE2 were administered 100 µg of ACE2.1mb protein by i.p. or i.v. injection or i.n. instillation. One day later, the mice were challenged with a high dose of SARS-CoV-2 USA-WA1/2020. For comparison, the mice were treated in parallel with the REGN-CoV2 cocktail, a mixture of REGN10933 and REGN10987 that has been shown to potently suppress SARS-CoV-2 replication in animal models (Baum et al., 2020). Viral RNA in the lung was quantified 3 days post-infection (dpi), the day of at which virus loads peak (Bao et al., 2020). The decoy strongly suppressed virus replication in the mice when administered either i.v. or i.n. Injection of the decoy i.p decreased the virus load 108-fold compared to mock while i.v injection decreased the virus load 15,700-fold and i.n. instillation decreased the virus load 26,500-fold, a level at which the viral RNA could not be undetected (**Figure 3A and B**). The monoclonal antibody cocktail closely mirrored the effect of the decoy protein in the three routes of administration. To compare the effectiveness of the sACE2, ACE2.mb, ACE2.1mb decoys, the proteins were injected i.v. and the mice were challenged with live virus. The results showed that sACE2 was the least effective while the ACE2.1mb decoy caused the greatest decrease in virus load (**Figure 3C**). Histological examination of the lung tissue of the mice showed a prominent infiltration of immune cells in the untreated mice (**Figure 3D)** that was largely absent in mice treated with sACE2 and completely prevented by treatment with the ACE2.1mb decoy. To understand the kinetics with which the decoy protein suppressed virus replication, mice were treated and then infected 24 hours later and viral RNA was measured every day over the course of one week. The results showed the absence of detectable viral RNA in the lung over the time course except for a small blip at 3-dpi; virus was not detected in the trachea over the time course (**Figure 3E**). A dose-response analysis of the potency of the decoy administered i.v. and i.n. showed that 100 μg of the protein suppressed virus replication to undetectable levels and that as little as 10 μg suppressed virus replication 12-fold **(Supplementary Figure 2A)**; administration i.n. was slightly more effective than i.v. at the 50 and 10 microgram doses. A dose-response analysis of the REGN-COV2 cocktail showed that the potency of the decoy was similar to that of the cocktail administered i.v or i.n. **(Supplementary Figure 2B)**.

**Figure 3.**
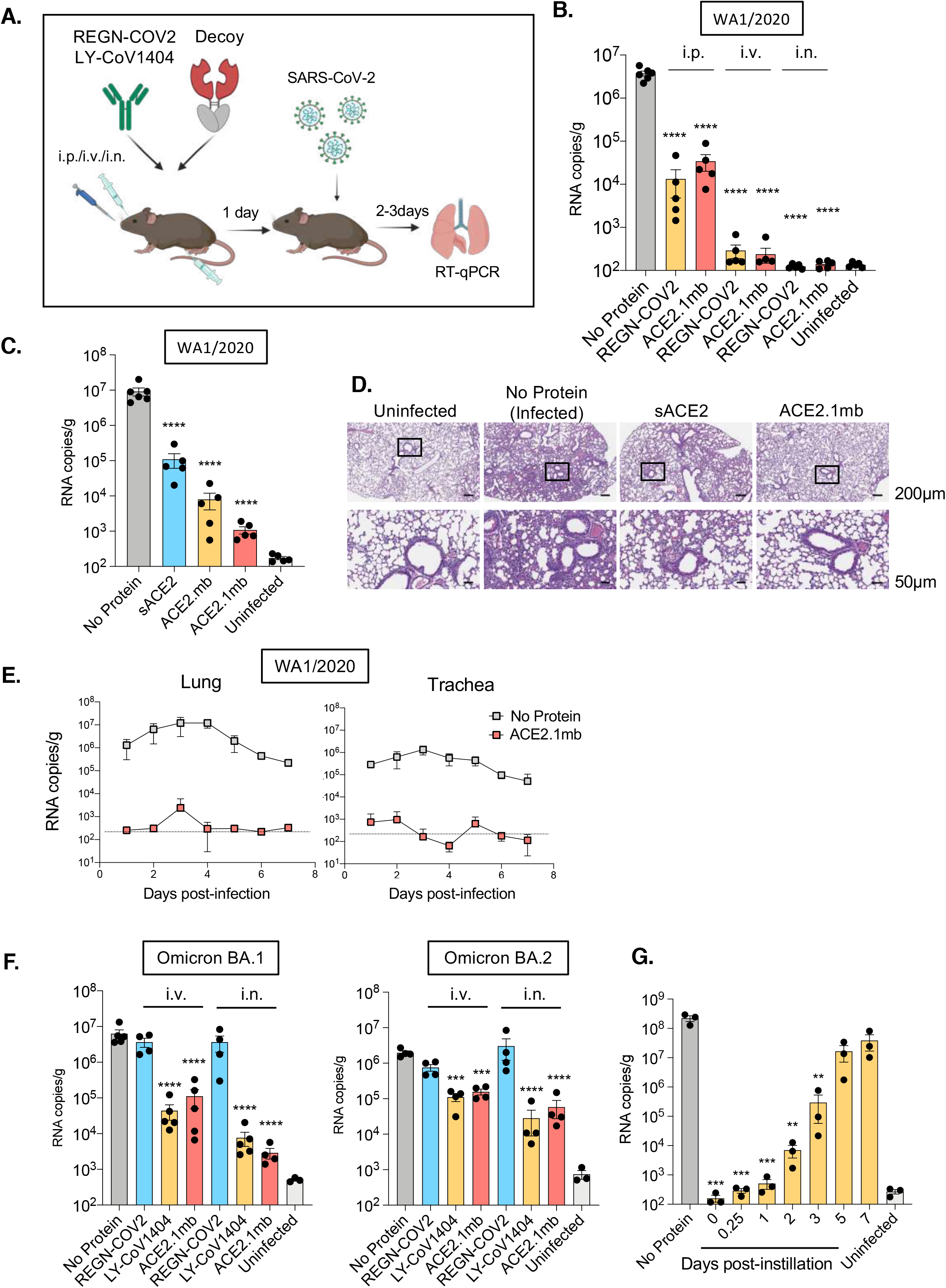
ACE2 decoy protects mice from SARS-CoV-2 infection and decreases virus loads. (A) Experimental scheme for decoy prophylaxis is shown. hACE2KI or BALB/c mice were injected i.p., i.v. or i.n. with 100 μg of ACE2.1mb or REGN-COV2 or LY-CoV1404. One day post injection, the mice were challenged with 2 × 10^4^ PFU of SARS-CoV-2 USA-WA1/2020 (hACE2KI) or SARS-CoV-2 Omicron BA.1 or BA.2 (BALB/c) and viral RNA copies were quantified 2-dpi (Omicron) or 3-dpi (WA1/2020). (B) Viral RNA copies (WA1/2020) in lung were quantified 3-dpi. (C) sACE2, ACE2.mb or ACE2.1mb proteins (100 μg) was administered to hACE2KI (n=5) by i.n. instillation. The following day, the mice were challenged with 2 × 10^4^ PFU of SARS-CoV-2 USA-WA1/2020. 3 dpi, subgenomic viral E RNA in the lung was quantified by RT-qPCR. (D) Hematoxylin and eosin (HE) staining of lung sections from SARS-CoV-2 WA1/2020 infected mice. Mice were injected with sACE2, ACE2.1mb. Post 1 day of injection, mice were challenged with SARS-CoV-2. 3-dpi, lung histology was visualized with HE staining. Scale bars, 200 μm (top), 50 μm (bottom). (E) ACE2.1mb protein (100 μg) was administered to hACE2KI mice (n=4) by i.n. instillation and the following day, challenged with 2 × 10^4^ PFU of SARS-CoV-2 USA-WA1/2020. At 1-, 2-, 3-, 4-, 5-, 6- and 7-dpi, subgenomic viral E gene RNA in the lung and trachea were quantified. The horizontal line indicates the level of detection determined using uninfected mouse tissues. (F) BALB/c mice were injected i.v. or i.n. with 100 μg of ACE2.1mb or LY-CoV1404 antibody (n=4-5). One day post injection, the mice were challenged with 2 × 10^4^ PFU Omicron BA.1 (left) or BA.2 (right). 2-dpi, lung subgenomic viral E RNA was quantified by RT-PCR. The copy numbers detected in uninfected samples is the result of low-level background priming. Confidence intervals are shown as the mean ± SD. **P ≤ 0.01, ***P≤0.001, ****P≤0.0001. The experiment was done twice with similar results. (G) hACE2KI mice were instilled i.n. with 100 μg of decoy proteins and then infected 0, 0.25, 1, 3, 5, and 7 days later with SARS-CoV-2 WA1/2020. At 3-dpi, lung subgenomic viral E RNA was quantified by RT-qPCR.

The efficacy of the decoy against the Omicron variants was tested in BALB/c mice which support high levels of replication of the virus (Halfmann et al., 2022). As controls, the mice were treated with the LY-CoV1404 monoclonal antibody which is active against the Omicron variant (Iketani et al., 2022; Liu et al., 2022; Tada et al., 2022a) or with the REGN-COV2 cocktail which is inactive against Omicron (Cameroni et al., 2021; Cao et al., 2021; Hoffmann et al., 2021; Iketani et al., 2022; Liu et al., 2022; Planas et al., 2021a; Tada et al., 2022a; VanBlargan et al., 2022). The following day, the mice were challenged with BA.1 or BA.2 virus. The decoy decreased the BA.1 virus load 56-fold by i.v. injection and BA.2,100-fold by i.n. instillation. The decreases were comparable to that caused by LY-CoV1404 (**Figure 3F, left**). The REGN-COV2 cocktail had no effect. The decoy was also active against BA.2. The decrease in virus load was not as dramatic but was similar to that of the highly potent LY-CoV1404 at the same dose (**Figure 3F, right and Supplementary Figure 2B)**. The durability of protection was analyzed by administering the decoy at increasing times pre-infection. The results showed a high degree of protection when the decoy was administered up to 2 days prior to infection and a moderate degree of protection 3-dpi (200-fold decrease in virus load); the protection was lost 5-dpi (**Figure 3G)**.

### Decoy suppresses virus load post-infection

To determine whether the decoy could be used to treat an established infection, hACE2KI mice were infected with USA-WA1/2020 and then treated 1-, 6-, 12-and 16-hours later injected i.p., i.v. or instilled i.n. with the ACE2.1mb decoy or REGN-COV2 cocktail (**Figure 4A**). At 3-dpi, virus loads in the lung were measured. I.p. injection of the decoy or monoclonal antibody had no effect on virus load at any of the time points (**Figure 4B**). In contrast, 1-hour post-infection, the i.v. injected decoy decreased the virus load 1,100-fold while i.n. instillation decreased the virus load 27,500-fold to an undetectable level. Virus loads were similarly decreased when the decoy was administered 6 hours post-infection. At 12 hours post-infection, the i.v. injected decoy had no significant effect while the i.n. instilled decoy decreased the virus load 20-fold, an effect that was maintained at the 16-hour time point. The effect of the decoy was comparable to that of the REGN-COV2 cocktail at all time points. The effect of the decoy was not as pronounced on Omicron BA.1 or BA.2 with decreases of 290-fold and 30-fold, respectively for the i.n. instilled decoy (**Figure 4C**). The effect of the decoy was 4-50-fold higher than LY-CoV1404 (**Figure 4C**).

**Figure 4.**
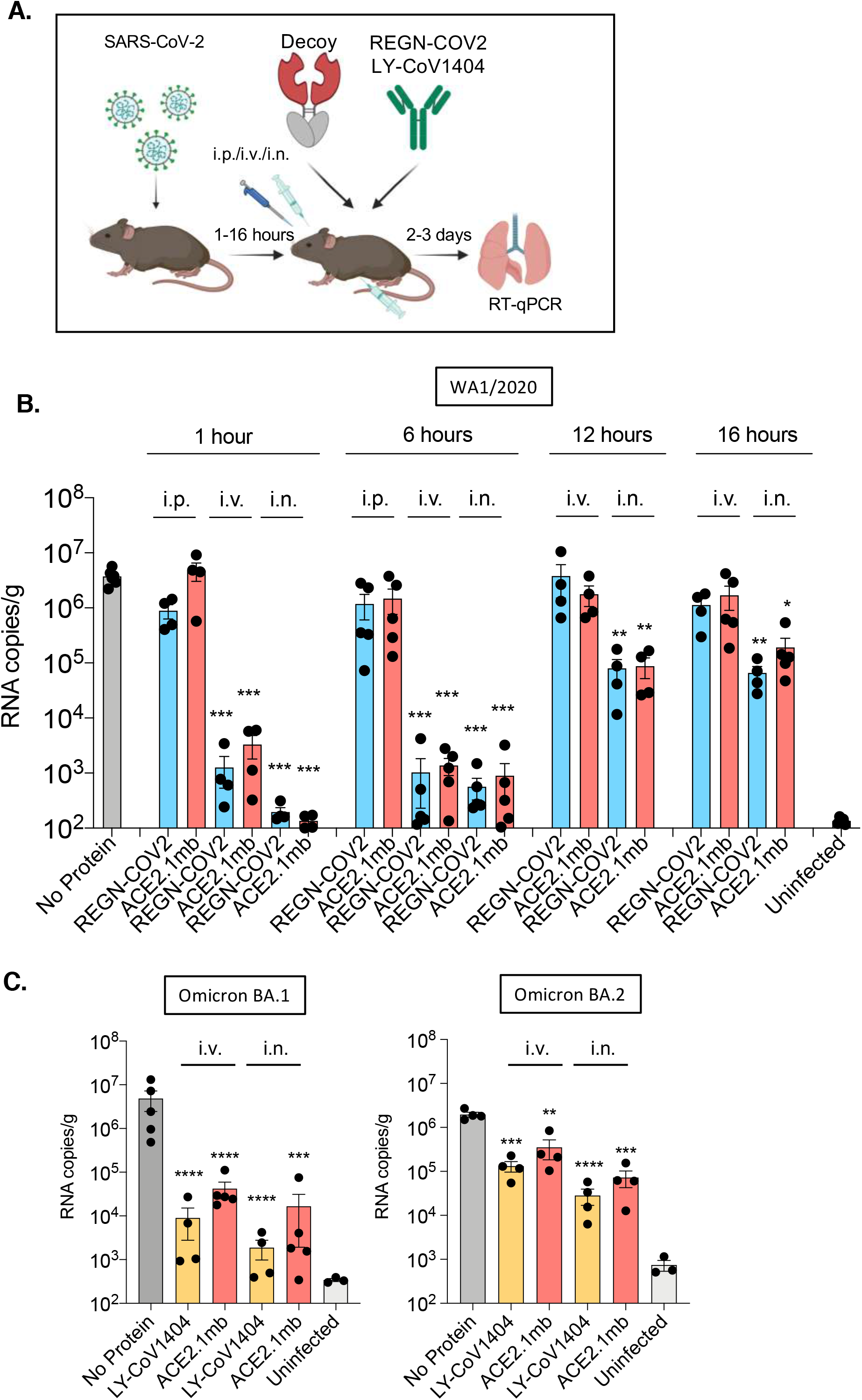
ACE2 decoy treats SARS-CoV-2 infection and decreases virus loads in mice. (A) Therapeutic experimental scheme for decoy treatment is shown. hACE2KI mice were infected with USA-WA1/2020 virus (n=4-5) and BALB/c mice were infected with Omicron BA.1 or BA.2 virus via i.n. injection. After 1, 6, 12, 16 hours post infection, mice were treated with 100μg ACE2.1mb or REGN-COV2 or LY-CoV1404 via i.p., i.v. or i.n. injection. Viral RNA copies in lung were quantified 2-dpi (Omicron) or 3-dpi (WA1/2020). The copy numbers detected in uninfected samples is the result of low-level background priming. (B) After 1, 6, 12, 16 hours post infection, mice were treated with decoy. Viral RNA copies (WA1/2020) in lung were quantified -dpi. (C) After 6 hours post infection, mice were treated with decoy. Viral RNA copies (Omicron BA.1 (left) or BA.2 (right)) in lung were quantified 2-dpi. Confidence intervals are shown as the mean ± SD. **P ≤ 0.01, ***P≤0.001, ****P≤0.0001. The experiment was done twice with similar results.

## Discussion

We report here that a high affinity ACE2 microbody decoy was highly effective both in the form of a recombinant protein for the treatment of SARS-CoV-2 infection. The decoy was highly potent *in vitro* against viruses with Variants of Concern spike proteins including Omicrons BA.1 and BA.2. I.v. injection or i.n. instillation of the recombinant decoy protein prior to infection protected hACE2KI mice from a high dose of live virus, suppressing virus replication to undetectable levels and preventing lung pathology. The decoy was highly effective administered up to 3 days prior to infection. Administration of the recombinant protein shortly after infection with SARS-CoV-2 rapidly suppressed virus replication in the lung. The decoy was at least as potent for prophylaxis and treatment as potent emergency-use-authorized monoclonal antibodies. The addition of the single CH3 IgG1 domain served to extend the half-life of the decoy and increased its avidity for the spike protein while preventing absorbance of the protein to Fc receptors that would decrease the concentration of the free protein.

This report confirms and extends recent findings from other groups using decoys containing full-length Fc regions that were reported during the preparation of this manuscript. Hoshino’s group reported that an ACE2.Fc fusion protein containing mutations A25V, K31N and N90H administered 2 hours post-infection increased the survival in the hamster model (Higuchi et al., 2021; Ikemura et al., 2022). Zhang *et al*. reported that a high affinity receptor decoy termed sACE2v2.4.Fc fusion protein containing mutations T27Y, L79T, and N330Y (Chan et al., 2020) protected K18-ACE2 transgenic mice from infection with SARS-CoV-2 variants and protected mice from disease when given 12- and 24-hours post-infection (Zhang et al., 2022).

I.v. injection or i.n. instillation of the decoy protein 1-or 6-hours after infection with USA-WA1/2020 decreased the virus load 10,000-fold, demonstrating the potency with which it suppresses virus replication. Administration at later times (12- and 16-hours post-infection) was less effective, decreasing virus loads by 100-fold. This timing should not be taken to mean that in humans decoy therapy would need to be administered as soon post-infection. In the mouse model, virus replication kinetics are somewhat faster than in humans, peaking 2-4-dpi, due to the high dose of virus administered (Jones et al., 2021). In our study, the effect of the decoy was similar to that of the therapeutic monoclonal antibodies which are effective at preventing hospitalization and death when given during the symptomatic phase of infection, several dpi (Group et al., 2021; Razonable et al., 2021). The decoy may act in humans with kinetics similar to that of anti-spike protein monoclonal antibodies that neutralize virus by a similar mechanism. Results reported here with the recombinant protein are consistent with those previously reported with a high affinity decoy 3N39v2 containing 4 mutations (Higuchi et al., 2021; Ikemura et al., 2022) and with the decoy sACE2v2.4-Ig that showed protection against infection and disease in hamsters and transgenic mouse models.

The antiviral effect of the decoy was influenced by its route of administration. I.p. injection localized the protein mainly to the liver, serum and spleen. This route of decoy administration was not effective as was the case for i.p. injected monoclonal antibody. I.v. injection resulted in a higher concentration of the decoy in the lung and potent antiviral activity. I.n. instillation localized the decoy primarily to the lung and trachea, as might be expected, and provided in the highest degree of protection. The findings suggest that the decoy acts in the lung to suppress virus replication and that both i.v and i.n. are effective routes of administration.

The decoy was less protective against the BA.1 and BA.2 subvariants both *in vitro* and *in vivo* than against the parental USA-WA1/2020 virus. Its activity against the variants was comparable to that of the potent therapeutic monoclonal antibody LY-CoV1404 suggesting that the decoy would be similarly effective against these variants in clinical use. The decoy has increased potency against the recent BA.2.75 (data not shown), suggesting that the virus is not mutating in such a way as to decrease effectiveness of the decoy approach.

The rapid evolution of SARS-CoV-2 has presented an obstacle to the development of effective therapies that target the spike protein. *In vitro* selection studies suggest that the spike protein will continue to evolve over the next several years imposing further challenge to the development of broadly neutralizing monoclonal antibodies (Schmidt et al., 2021). The receptor decoy approach is more resistant to immuno-evasion by novel variants because of the requirement that the spike protein preserves its affinity for ACE2. As new spike protein variants evolve, they are likely to remain susceptible to neutralization by the decoy. As the virus has continued to evolve and increase its transmissibility following zoonosis into humans, the variant spike proteins have tended to increase their affinity for ACE2, resulting in increased sensitivity to neutralization by the decoy (Cao et al., 2022a; Yue et al., 2023). It is conceivable that a variant will emerge that switches its receptor usage to an alternative cell surface protein, thereby becoming resistant to the decoy; however, such an event has not occurred in nature and extensive laboratory mutagenesis of the spike protein has not resulted in a receptor switch (Greaney et al., 2021; Starr et al., 2020). If such a switch were to occur in a pandemic coronavirus or virus of another virus class, a receptor decoy could be developed based on its receptor and rapidly deployed.

### Limitations of Study

The study is based on analyses of decoy receptor in hACE2KI mice. While these mice express human ACE2 at physiological levels in the appropriate tissues, they may not entirely accurately reflect the human. Calculations of how much protein would be required to treat humans may not be accurate. Moreover, the time course of SARS-CoV-2 infection is longer in humans, and thus the time courses analyzed here may not be directly translatable to human.

## Acknowledgements

We thank Victor Torres and Juliana Ilmain (NYU Grossman School of Medicine) for assistance with FPLC purification of recombinant proteins and Meike Dittman and Bruno Rodriguez-Rodriguez for ACE2.TMPRSS2.Vero E6 cells. We thank members of the Experimental Pathology Research Laboratory which is supported by NIH Grant P30CA016087 to the NYU Langone Laura and Isaac Perlmutter Cancer Center for histology. The work was funded by grants from the NIH to N.R.L. (DA046100, AI122390 and AI120898). T.T. was supported by the Vilcek/Goldfarb Fellowship Endowment Fund.

## Author contributions

T.T., B.M.D. and H.Z. conducted the experiments. T.T. designed the experiments and wrote the paper. T.T. and B.M.D. did the statistical analysis. N.R.L. supervised the study and revised the manuscript.

## STAR Methods

### Resource Availability

#### Lead Contact

Further information and requests for resources and reagents should be directed to and will be fulfilled by the Lead Contact, Nathaniel R. Landau.

#### Materials Availability

All unique DNA constructs, proteins and pseudotyped virus generated in this study are available from the Lead Contact upon request.

#### Data and Code Availability

- The data used in this study are available upon request from the lead contact.
- This paper does not report original code.
- Any additional information required to reanalyze the data reported in this paper is available from the lead contact upon request.

##### Experimental Model and Subject Details Mice

hACE2KI (B6.129S2(Cg)ACE2tm1(ACE2)Dwnt/J) and BALB/c mice were purchased from the Jackson Laboratory. We collected as many mice as possible and used them for the experiments. The sample size was described in the Figure legend. All animal experiments were done under protocols approved by the NYU Langone Institutional Animal Care and Use Committee (#170304) in accord with the standards set by the Animal Welfare Act. The study was approved by the NYU School of Medicine Division of Comparative Medicine Standard Operating Protocol (40-008-17). All experiments were done twice or triplicates with similar results.

##### Cells

293T and Vero E6 cells were cultured at 37°C under 5% CO_2_ in Dulbecco’s modified Eagle medium (DMEM) supplemented with 10% fetal bovine serum (FBS) and penicillin/streptomycin. ACE2.293T(Tada et al., 2020) and ACE2.TMPRSS2.Vero E6 cells were cultured with the addition of puromycin (1 μg/ml). ExpiCHO-S cells (Thermo Fisher Scientific) were grown at 37 °C under 8% CO_2_ in suspension in ExpiCHO serum-free expression medium in a shaking incubator.

##### Plasmids

The plasmids pLenti.GFP.nLuc, pMDL, pcRev, pcCOV2.S.delta19 and pcCOV2.S.delta19-variant spikes used to produce spike protein-pseudotyped lentiviruses have been previously described (Tada et al., 2020). The decoy expression vectors pcsACE2, pcACE2mb and pcACE2.H345A.mb have been previously described (Tada et al., 2020). To construct the expression vector pcACE2.1mb encoding the high affinity microbody decoy ACE2.1mb, point mutations T27Y, L79T, N330Y were introduced into pcACE2.H345A.mb by overlap extension PCR and the amplicon was cloned into the Kpn-I and Xho-I sites of pcDNA6. To construct the expression vector pcsACE2-nLuc and pcsACE2.1mb-nLuc expressing a decoy:nanoluciferase protein, DNA fragments encoding ACE2 amino acids 1-741 and nanoluciferase were amplified by PCR and joined by overlap extension PCR using primers containing Kpn-I and Xho-I sites. The resulting amplicon was cloned into Kpn-I and Xho-I cleaved pcDNA6. Nucleotide sequences of all plasmids were confirmed by DNA sequencing.

##### Monoclonal antibodies

REGN-COV2 (REGN10933+REGN10987) were provided by Regeneron Pharmaceuticals. Bebletovimab (LY-CoV1404) was obtained from discarded vials.

##### Recombinant protein purification

ExpiCHO-S cells (Thermo Fisher Scientific) cultured in shaker flasks in serum-free medium were grown to a density of 6 × 10^6^/ml and transfected with 400 μg of plasmid DNA using with 1.28 ml of ExpiFectamine. After 12 hours, 2.4 ml of ExpiCHO Enhancer and 64 ml of ExpiCHO Feed were added. After 4 days, the culture supernatant was collected and passed over a 0.22 μm filter. The supernatant was passed over a 5 ml HiTrap Chelating column charged with nickel on an Akta FPLC (GE healthcare). The column was washed with buffer containing 20 mM Tris pH 8, 150 mM NaCl, 10mM imidazole and the bound protein was then eluted in buffer containing 250 mM imidazole. The eluate was loaded onto a Superdex 200 size-exclusion column (GE healthcare) in running buffer containing 10 mM Tris pH 7.4, 150 mM NaCl. Fractions were collected and those containing peak protein concentrations were pooled. Protein purity was analyzed on a 4-12% Bis-Tris SDS-PAGE by silver staining (Invitrogen).

## Method Details

### Virion-decoy pull-down assay

Decoy proteins (5, 2, 0.5 and 0.1 μg) were allowed to bind 30 μl of nickel-nitrilotriacetic acid-agarose beads (QIAGEN) for 1 hour after which unbound decoy was removed by washing with PBS. The beads were then incubated with 30 μl (30 μg) of D614G spike protein-pseudotyped virus for 1 hour after which unbound virions were removed by washing with PBS. The bound virions were then eluted from the beads with Laemmle loading buffer containing reducing agent (Invitrogen) and analyzed on an immunoblot probed with anti-p24 monoclinal antibody AG3.0 (Creative Biolabs) and horseradish peroxidase (HRP)-conjugated goat anti-mouse IgG secondary antibody (Sigma-Aldrich). The signals were developed with Luminata Crescendo Western HRP Substrate (Millipore) and membranes were visualized on an iBright imaging system (Invitrogen) (Tada et al., 2020).

### Cell-based decoy spike binding assay

293T cells (2 × 10^6^) were transfected with 2 μg spike expression vector by lipofection using lipofectamine 2000 (Invitrogen). One day post-transfection, the cells were plated in a 96 well plate at 1 × 10^4^ cells/well. The following day, decoy-nLuc fusion protein was added to the wells. After 30 minutes, the free fusion protein was removed by washing with PBS and cell-bound luciferase activity was measured using NanoGlo luciferase substrate (Nanolight) in an Envision 2103 microplate luminometer (PerkinElmer) (Tada et al., 2022b).

### Pseudotyped lentiviral neutralization assay

Spike protein-pseudotyped lentiviruses were generated as previously described (Tada et al., 2020). Briefly, virus stocks were generated by cotransfection of 293T cells with pMDL, pLenti.GFP.nLuc, pcCoV2.S-Δ19 and pRSV.Rev using the calcium phosphate method. After 2 days, the culture supernatant was harvested and the virus was concentrated by ultracentrifugation at 4°C for 1 hour at 30,000 X g and normalized for reverse transcriptase (RT) activity. To determine the neutralizing titer of the decoy proteins, serially diluted decoys were incubated with pseudotyped virus (MOI=0.2) for 30 minutes at room temperature and then added to ACE2.293T or ACE2.TMPRSS2.Vero E6 cells. At 2-dpi, luminescence was measured in an Envision 2103 microplate luminometer (PerkinElmer). All samples were assayed in duplicate and IC50s were calculate by Prism 8 software.

### Decoy localization *in vivo*

100 μg of ACE2.1mb-nLuc or sACE2-nLuc proteins were injected intraperitoneal (i.p.), intravenous (i.v.) or by intranasal (i.n.) instillation. After 6, 12, 24, 48, 72 hours, the mice were sacrificed and the tissues were homogenized in lysing matrix D tubes (MP Biomedicals) with a FastPrep-24 5G homogenizer (MP Biomedicals). Blood was collected by submandibular bleeding and serum was harvested. The tissue lysates were mixed with an equal volume of Nano-GLO Luciferase Assay Reagent (Nanolight) and luciferase activity was quantified on an Envision 2103 plate reader (PerkinElmer). Decoy concentration and half-life were determined based on a standard curve derived from the mixture of serial diluted ACE2.1mb-nLuc proteins and tissue lysate from wild-type. For live imaging of the decoy, mice were injected i.p. with 100 μl 1:40 diluted Nano-GLO substrate. After 3 minutes, the mice were live imaged on an IVIS Lumina III XR (PerkinElmer).

### Preparation of live virus

SARS-CoV-2 WA1/2020 P1 virus stock (BEI Resources, NR-52281) was amplified by a second round of replication on Vero E6 cells infected at MOI=0.01. At 3-dpi, the culture medium was harvested, filtered through a 0.45 μm filter and frozen at −80°C in aliquots. The virus was titered by plaque assay on Vero E6 cells (Wei et al., 2020). SARS-CoV-2 Omicron BA.1 (BEI Resources, NR56461) and BA.2 stocks (BEI Resources, NR-56781) were grown on ACE2.TMPRSS2.Vero E6 cells infected at an MOI=0.1. The P1 stock was amplified by a second round of replication on ACE2.TMPRSS2.Vero E6 cells infected at MOI=0.01. The virus-containing supernatant was filtered through a 0.45 μm filter, concentrated by passage through an Amicon filter (Millipore) and stored in aliquots at −80°C. The virus was titered by plaque assay on Vero E6 cells. Live virus was handled by trained personnel in a Biosafety level 3 facility.

### Prophylaxis and treatment of mice with decoy proteins

For prophylaxis, 6-8 weeks old hACE2KI or BALB/c mice were anesthetized with isoflurane and injected i.v. with decoy or monoclonal antibody, or alternatively, were anesthetized with ketamine– xylazine cocktail and administered the proteins by i.n. instillation. After 1 day, the mice were infected i.n. with 2 × 10^4^ PFU USA-WA1/2020 (hACE2KI) or Omicron BA.1 or Omicron BA.2 (BALB/c). At 2-dpi for Omicron-infected mice or 3-dpi for USA-WA1/2020-infected mice, the mice were sacrificed and lungs and trachea were harvested and homogenized. Littermate controls were included in all experiments. RNA was prepared from 200 µl of the lysate using the Quick-RNA MiniPrep kit (Zymo Research). For treatment experiments, hACE2KI mice were infected with 2 × 10^4^ PFU of SARS-CoV-2 USA-WA1/2020 i.n. At 1-, 6-, 12-or 16-hours post-infection, mice were administered therapeutic monoclonal antibodies or decoy protein (100 μg) i.p., i.v. or i.n. 3-dpi, lung and trachea were harvested and SARS-CoV-2 subgenomic E gene levels were quantified by RT-qPCR.

### RT-qPCR

Virus loads were measured by quantification of subgenomic viral E gene by RT-qPCR with TaqMan probes. Cellular RNA was mixed with TaqMan Fast Virus 1-step Master Mix (Applied Biosystems), 10 mM forward and reverse primers and 2 mM probe. PCR cycles were 95°C for 20s, 95°C for 3s, 40 cycles at 60°C for 30s) using forward primer E Sarbeco F (ACAGGTACGTTAATAGTTAATAGCGT), reverse primer E Sarbeco R (ATATTGCAGCAGTACGCACACA) and probe E Sarbeco P1(FAM-ACACTAGCCATCCTTACTGCGCTTCG-BHQ1) (Corman et al., 2020). E gene subgenomic RNA copies were measured using forward primer subgenomic F (CGATCTCTTGTAGATCTGTTCTC) (Emma S. Winkler, 2022), reverse primer E Sarbeco R and probe E Sarbeco P1). Absolute copy numbers were determined by normalization to a standard curve generated with *in vitro* transcribed synthetic RNA containing the E gene sequence (2019-nCoV_E Positive Control, IDT: 10006896). Cell lysate GAPDH copy numbers were measured as a control using mGAPDH.forward (CAATGTGTCCGTCGTGGATCT) and mGAPDH.reverse (GTCCTCAGTGTAGCCCAAGATG) with mGAPDH probe (CGTGCCGCCTGGAGAAACCTGCC). Data from tissue analyses was normalized to GAPDH. Virus load was determined by the 2−ΔΔCTmethod).

### Histology

Mice were infected with USA-WA1/2020 and sacrificed 3-dpi. Tissues were fixed in 10% buffered formalin and processed through graded ethanol and xylene solutions and then embedded in paraffin with a Leica Peloris automated processor. Five-micron sections were deparaffinized and stained with hematoxylin and eosin on a Leica ST5020 automated histochemical stainer. The slides were scanned at 40X on a Leica AT2 whole slide scanner.

### Data analysis and statistics

All experiments were in technical duplicates or triplicates. Statistical significance was determined by the two-tailed, unpaired t test using GraphPad Prism (Version 8) software. Significance was based on two-sided testing. Confidence intervals are shown as the mean ± SD. (*P≤ 0.05, **P≤ 0.01, ***P≤0.001, ****P≤0.0001).

---

**Supplementary Figure 1.**
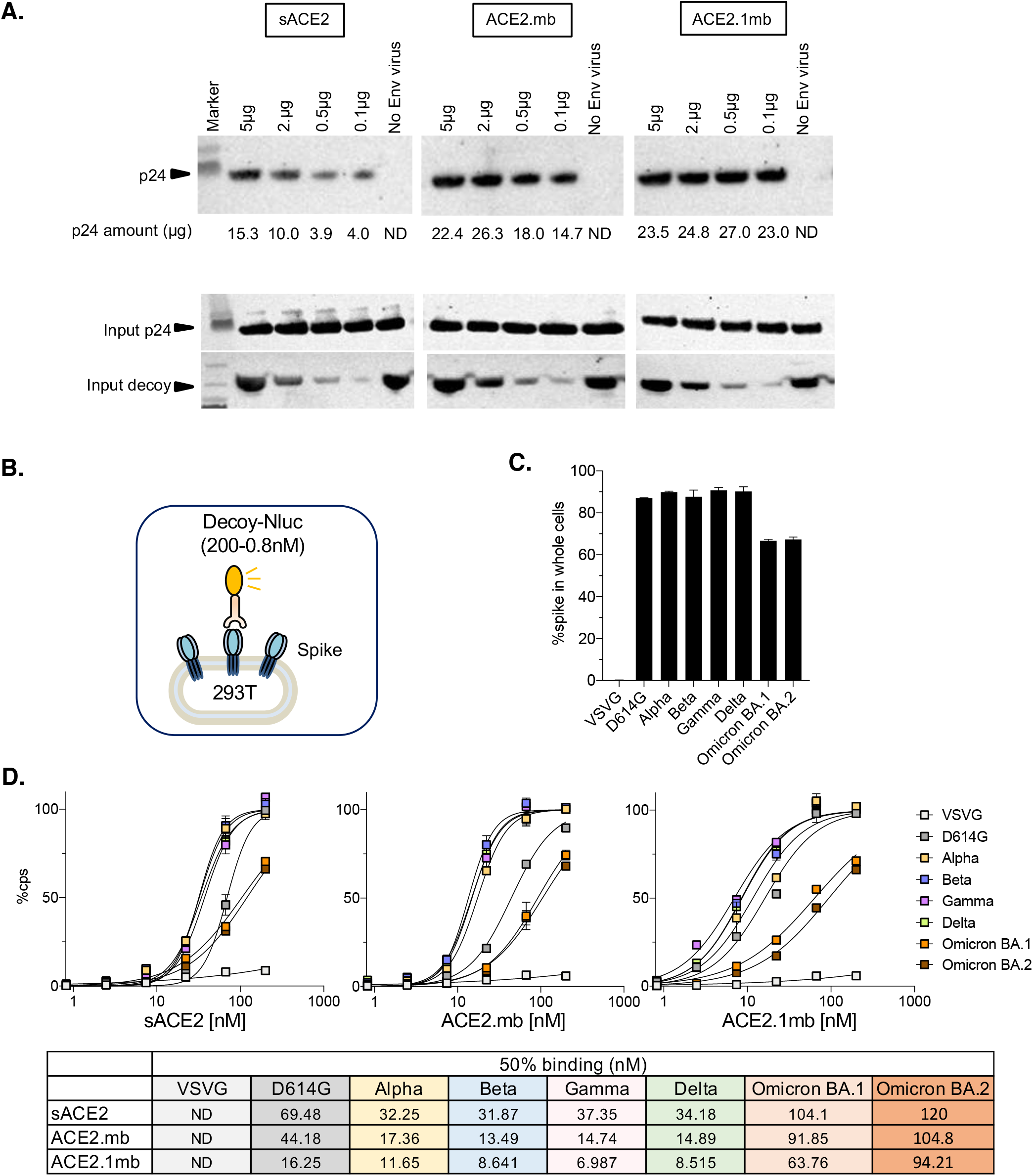
Increased affinity of ACE2.1mb for the spike protein. (A) Ni-NTA agarose beads were coated with different amounts (5, 2, 0.5, 0.1 μg) of sACE2, ACE2.mb or ACE2.1mb. A fixed amount of lentiviral virions pseudotyped with the D614G spike protein (30 μl) or control virions lacking the spike were incubated with the beads. After 1 hour, free virions were removed by centrifugation and the remaining bead-bound virions were detected on an immunoblot probed with anti-p24 antibody. The amount of bead-bound p24 capsid protein was calculated against a standard curve with recombinant capsid and is indicated below each lane. Input virions and decoy proteins were detected on an immunoblot probed with anti-p24 and anti-His antibody and are shown below. ND: Not detected. (B) The decoy:spike binding assay is diagrammed (left). 293T cells were transfected with 2 μg pcDNA-6 expression vectors for the VOC spike proteins. The cells were incubated at 37° with different amounts of decoy:luciferase fusion proteins. After 1 hour, unbound protein was removed by centrifugation and the amount of bound decoy protein was determined by luciferase assay. The amount of spike protein on the transfected 293T cells was analyzed by flow cytometry with anti-spike monoclonal antibody against the S2 protein (right). Avidity or the decoys is shown as curves with 100% binding set as luciferase activity at 400 nM decoy (below). The table shows the decoy concentration required for 50% maximal binding. The experiment was done three times with similar results.

**Supplementary Figure 2.**
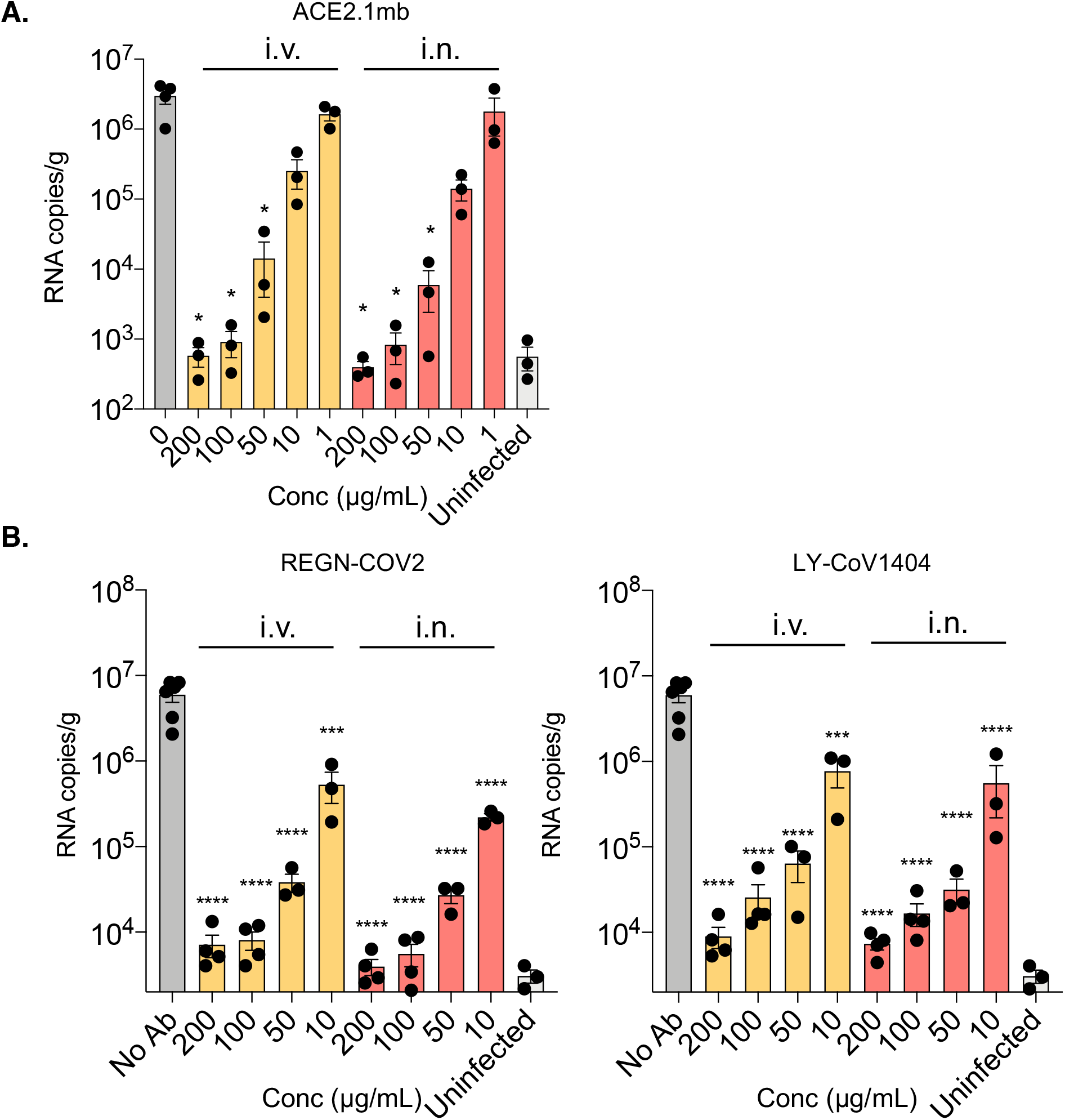
Decoy and therapeutic monoclonal antibodies protect mice from SARS-CoV-2 infection. (A) hACE2KI mice were injected i.v. or instilled i.n. with different amounts of decoy proteins. At 1-dpi, the mice were challenged with 2 × 10^4^ PFU SARS-CoV-2 USA-WA1/2020 (n=3). 3-dpi, lung subgenomic viral E RNA was quantified by RT-qPCR. (B) hACE2KI mice were injected i.v. or instilled i.n. with different amounts of REGN-COV2 or LY-CoV1404 antibody (n=3). At 1-dpi, the mice were challenged with 2 × 10^4^ PFU of SARS-CoV-2 USA-WA1/2020. At 3-dpi, lung subgenomic viral E RNA was quantified by RT-qPCR.

